# The ModelSEED Database for the integration of metabolic annotations and the reconstruction, comparison, and analysis of metabolic models for plants, fungi, and microbes

**DOI:** 10.1101/2020.03.31.018663

**Authors:** Samuel M. D. Seaver, Filipe Liu, Qizhi Zhang, James Jeffryes, José P. Faria, Janaka N. Edirisinghe, Michael Mundy, Nicholas Chia, Elad Noor, Moritz E. Beber, Aaron A. Best, Matthew DeJongh, Jeffrey A. Kimbrel, Patrik D’haeseleer, Erik Pearson, Shane Canon, Elisha M. Wood-Charlson, Robert W. Cottingham, Adam P. Arkin, Christopher S. Henry

## Abstract

For over ten years, ModelSEED has been a primary resource for the construction of draft genome-scale metabolic models based on annotated microbial or plant genomes. Now being released, the biochemistry database serves as the foundation of biochemical data underlying ModelSEED and KBase. The biochemistry database embodies several properties that, taken together, distinguish it from other published biochemistry resources by: (i) including compartmentalization, transport reactions, charged molecules and proton balancing on reactions;; (ii) being extensible by the user community, with all data stored in GitHub; and (iii) design as a biochemical “Rosetta Stone” to facilitate comparison and integration of annotations from many different tools and databases. The database was constructed by combining chemical data from many resources, applying standard transformations, identifying redundancies, and computing thermodynamic properties. The ModelSEED biochemistry is continually tested using flux balance analysis to ensure the biochemical network is modeling-ready and capable of simulating diverse phenotypes. Ontologies can be designed to aid in comparing and reconciling metabolic reconstructions that differ in how they represent various metabolic pathways. ModelSEED now includes 33,978 compounds and 36,645 reactions, available as a set of extensible files on GitHub, and available to search at https://modelseed.org and KBase.

## INTRODUCTION

Genome-scale metabolic reconstructions and models have become central tools for systems biology research. These models are valuable for their capacity to consolidate and represent the functional annotations of biology using the more concrete and universal language of biochemistry. By representing annotations with chemistry, we can move beyond a simple cataloguing of observed and predicted functions, and begin to assemble those functions into the interconnected series of metabolic pathways that comprise the chemical foundation of any metabolic model. Models can then be applied to automatically identify any gaps that interrupt these pathways and suggest new hypothesis-driven experiments to fill these gaps (1).

Beyond the capacity of models to give structure and chemical meaning to functional annotations in biology, these models are also valuable for their predictive capacity. Today, models can be used to predict a wide range of biological phenotypes, including: (i) respiration, photosynthesis, and fermentation types (2–11); (ii) feasible growth conditions and Biolog phenotype array profiles (12–15); (iii) essential genes and reactions (16–20); (iv) potential existing or engineerable by-product biosynthesis pathways (21–25); and (v) the yields and even titre available for those pathways (26–29).

Metabolic models are also now emerging as ideal tools for the integration of fluxomes, metabolomes, transcriptomes, and proteomes. This capability has been applied to empower the development and parameterization of dynamic kinetic models (30–32), the reconstruction of tissue specific metabolic models (33–35), the discovery of new chemistry and pathways from metabolomes (36), and the simulation and analysis of interactions within a microbial community (37–39).

Given the rapid adoption of these models as tools in systems biology, the pace with which new models are produced has grown dramatically, particularly with the emergence of numerous automated model reconstruction pipelines (40). The diversity of resources now producing large numbers of these models has created new challenges due to a lack of standardization in models and their underlying biochemistry, assumptions, and associated data. Tools like MEMOTE aid in improving standardization in metabolic models (41), but variations in how the same metabolic pathways are represented in different models remains a problem. The ability to rapidly map chemistry across different metabolic models is critical to facilitate the comparison and reconciliation of models or to permit models to interoperate within larger microbiome community models (42). It is also critical to support the integration of supplementary data for models, including thermodynamic properties (43), kinetic constants (44), and metabolomics data (45, 46).

Here we present the ModelSEED biochemistry database, a transparent resource of biochemistry designed to support standardization and data integration. This database embodies several properties tailored to this objective. First, biochemistry data are unified and integrated from multiple major external sources, including KEGG (47, 48), MetaCyc (49), and BiGG (50). All reactions and compounds from these sources are integrated and retained within the database to facilitate rapid automated mapping of new models to the database. Second, special attention and curation are performed to ensure that as many compounds in the database as possible have chemical structures associated with them, and compounds with identical structures are mapped together within the database. This facilitates the checking of reaction mass and charge balance, and the mapping of database metabolites to metabolomics data. Third, thermodynamic properties and pH-based molecular ion charges are computed consistently for compounds in the database, with these data being further used to compute reaction properties, including proton stoichiometry, Gibbs energy change of reaction, and predicted reversibility and directionality. These reaction and compound properties may then be mapped to models, where they can be used to evaluate thermodynamic feasibility of model output. Fourth, we applied flux balance analysis to explore how the connectivity of this new release of ModelSEED has improved in terms of activating diverse pathways and simulating biomass production in diverse media. This new release of the ModelSEED database also includes an ontology, which maps equivalent reactions from various data sources to each other. This ontology can be used to automatically convert a model to a standard biochemical representation to facilitate rapid comparison and integration. Finally, the database is encoded within GitHub, with a collection of testing scripts and a continuous integration environment, designed to facilitate the rapid extension of the ModelSEED database with community contributions, as well as providing an update and release mechanism enabling users to sync with database changes and see full details on how the database changes with each update cycle. Such extensibility is critical to keep pace with rapid discovery of new chemistry. Below we describe each of these capabilities in detail.

## MATERIAL AND METHODS

### Gathering and integration of biochemical data

We downloaded the molecular structures from KEGG and MetaCyc, and the compounds and reactions from >20 biochemistry databases and published metabolic models. See Supplemental Table 1 for a full list of sources.

### Protonation and conversion of ModelSEED compounds

Marvin from ChemAxon was used to protonate all molecular structures at a pH of 7 and to convert every molecular structure into InChI and SMILES format, Marvin 19.1, ChemAxon (https://www.chemaxon.com). Due to limitations in (i) the molecular structures; (ii) the InChI format; and (iii) Marvin, we are unable to protonate or convert every molecular structure to InChI or SMILES format. Where possible, we defaulted to the InChI representation of the protonated structure and, failing that, the SMILES representation of the unprotonated structure.

### Biochemical integration

We successively integrated the downloaded biochemistry in multiple layers (Table 1 and Figure 1), prioritizing first the KEGG and MetaCyc biochemistry as primary sources containing molecular structures, then selected BioCyc databases and published models as secondary sources that use KEGG/MetaCyc identifiers. The rest of the published models were tertiary sources, and finally Rhea and MetaNetX using external identifiers. At each stage, we integrated the compounds first, then integrated reactions based on whether they use the same reactants, products, and stoichiometry (allowing for variations in proton stoichiometry). Crucially this means we did not integrate reactions based on names or identifiers. For the integration of the primary sources, we used InChI and SMILES representations of the available molecular structures to match compounds from KEGG and MetaCyc. For the primary and secondary sources, if a new compound did not have an available structure, or a matching KEGG/MetaCyc identifier, we used the available synonyms to find matches if any.

**Figure 1.**
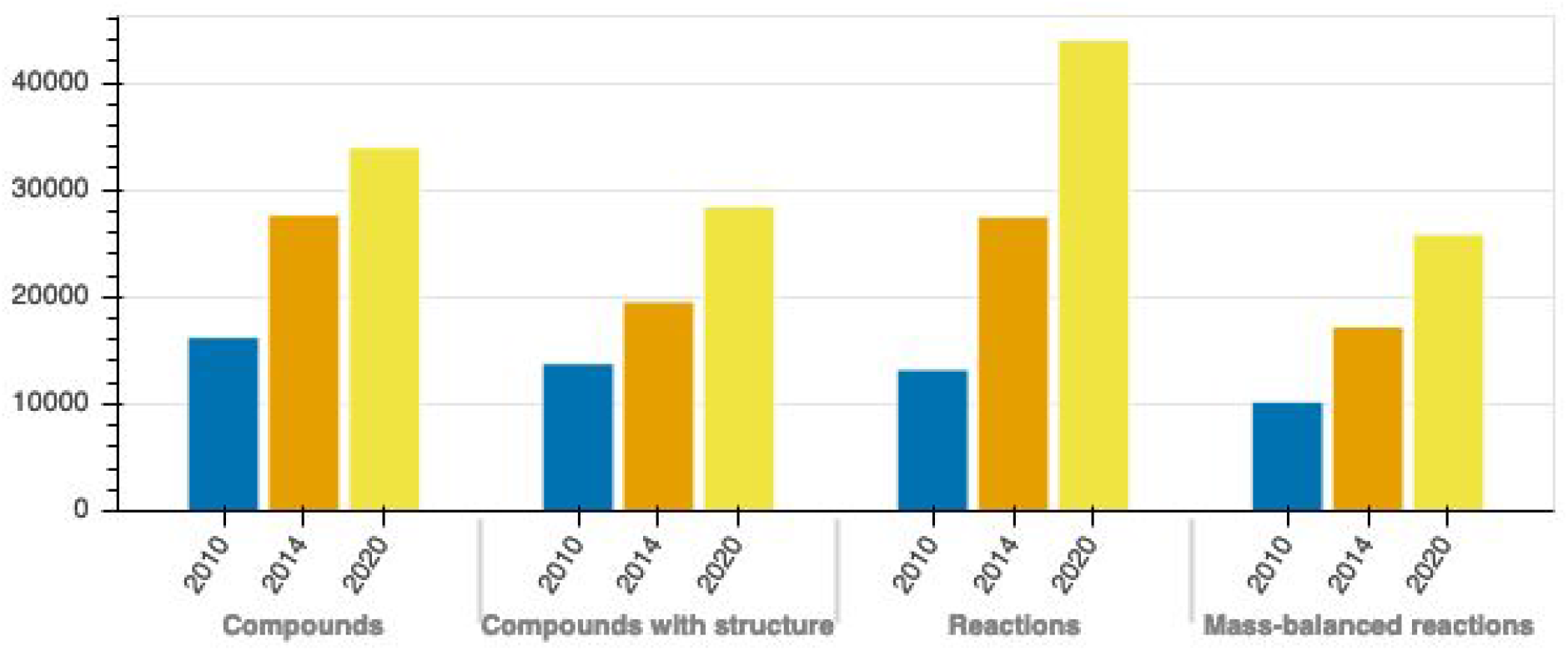
The growth of the ModelSEED biochemistry database. Since the release of the ModelSEED resource, along with its biochemistry, we have steadily updated the biochemistry database with the latest data in several public databases as well as integrated more published metabolic reconstructions. At the same time, we’ve refined our approach for integrating structural data, and so our database has not only grown in size, but also in quality: today we have a biochemistry database of more than 20,000 mass-balanced reactions that can be utilized in metabolic reconstructions spanning the microbial, fungal, and plant kingdoms.

**Table 1.** The degree to which ModelSEED biochemistry was integrated from different data sources (structure/identifier/synonym). It should be noted that the integration was done in stages, so “unintegrated” for KEGG and MetaCyc means not integrated with each other. “Unintegrated” for published models means not integrated with either KEGG, MetaCyc, or BioCyc. For KEGG/MetaCyc, the number in parentheses are for integration using structures only, and for BioCyc, the numbers in parentheses are for integration using BioCyc identifiers only.

| Sources | Integrated Compounds | Integrated Reactions | Unintegrated Compounds | Unintegrated Reactions |
| --- | --- | --- | --- | --- |
| KEGG/MetaCyc | 6,276 (6,219) | 4,791 (4,679) | 24,352 (22,268) | 23,376 (19,987) |
| BioCyc<br>(11 databases*) | 6,957 (6,893) | 6,724 (6,553) | 997 | 1,626 |
| Published models<br>(34 models) | 3,694 | 4,359 | 1,792 | 5,464 |

### Balancing of reactions in the ModelSEED

A combination of RDKit 2020.03.1.0 (https://www.rdkit.org; 51) and OpenBabel 2.4.1 (https://openbabel.org; 52), two open source cheminformatics software packages, were used to derive the correct formula and charge from the molecular structures. Having assigned the formula and charges of the protonated string, we calculate the mass and charge balance of every reaction in the database.

### Computation of thermodynamic properties of ModelSEED compounds and reactions

For previous releases of the ModelSEED biochemistry, the standard Gibbs energy of formation for each compound (**Δ**_f_G’^o^) and the standard Gibbs energy of reaction for each reaction (**Δ**_f_G’^o^) was estimated using a group contribution approach (53). For this new release, we re-calculated these energies using eQuilibrator, a more recent approach developed by Noor *et al*., 0.2.5 (http://equilibrator.weizmann.ac.il)(54). All energies were calculated at pH 7.0, ionic strength of 0.25 M, and temperature of 298.15 K. Data from these two methods were integrated in a complementary manner, giving precedence to the results from eQuilibrator.

The set of complete structures in the ModelSEED biochemistry from which energy of formation was computed using the group contribution approach, as integrated from KEGG and MetaCyc, only partially overlaps with the set of complete structures in MetaNetX, from which eQuilibrator computed pKa values. There were a total of 24,081 unique InChI structures in ModelSEED, and 465,752 unique InChI structures in MetaNetX, but only 19,520 of these structures were shared between the two databases. In addition, neither the group contribution method nor eQuilibrator was able to return an estimate for (**Δ**_f_G’^o^) for every structure, and as such, there were only 19,761 ModelSEED compounds with an estimate for (**Δ**_f_G’^o^) from the group contribution method and 17,602 ModelSEED compounds with an estimate for (**Δ**_f_G’^o^) from eQuilibrator. For each compound, we use the value from eQuilibrator in our database where possible, but we retain the values computed by the group contribution approach in the repository.

A reaction was considered to be “complete” when every reactant had a defined structure and for which (**Δ**_f_G’^o^) was available via either the group contribution method or eQuilibrator. There were 19,486 ModelSEED reactions defined as complete by the group contribution method, and 17,763 ModelSEED reactions defined as complete by eQuilibrator, with 15,106 reactions shared between the two. For each of these reactions, we used the value of (**Δ**_f_G’^o^) computed by eQuilibrator, except when estimated error returned by eQuilibrator exceeds an arbitrary value of 100 kilocal.mole^-1^ (2,503 reactions). For those, the value computed by the group contribution method was used. When computing **Δ**_f_G’^o^, we always used either eQuilibrator or group contribution values exclusively, and never mixed and matched **Δ**_f_G’^o^ values from these two methods to compute a single **Δ**_f_G’^o^ because they differ in their reference states, and thus would cause significant error. For each of the reactions with an estimated **Δ**_f_G’^o^, we applied a heuristic to estimate the thermodynamic reversibility of the reaction based on a set of rules developed in earlier work (55).

### Undetermined compounds

Many compounds in the database, whether they were assigned a structure or not, were considered to be undetermined for a number of reasons. The compound may be “lumped” if their structure was partially or wholly unknown and the “lumped” structure was represented by an “R” group. Where we could, we included the reactions containing such compounds while making sure that the “R” groups balanced and represented the same sub-structure. The compound may have been generic or abstract, in that they were representative of a class of compounds; so we included these compounds by generating hierarchical links between them and their structurally-specific representatives.

### GitHub policies for community contributions

GitHub offers a valuable venue and toolkit to support community curation, and it has been applied to this purpose for the co-development of computer code by a vast user community. This community has demonstrated how a large group of individuals can work together on a single project and be effective. The tools that GitHub offers its developer-users, as well as the policies and practices it encourages, are critical components that make large-scale cooperative projects possible. With this release of the ModelSEED biochemistry database, we anticipate the same principles and methods apply and will ultimately support large-scale community-curation of biochemistry data in the ModelSEED. To accomplish this goal, we have adopted many of the same practices used by developers.

#### Use of branches

The ModelSEED repository includes a dev and master branch in GitHub. All releases will be deployed to the master branch, which will be tagged with release identifiers when the release is complete. All active new curation work will take place in the dev branch, where all external contributors are encouraged to submit their pull requests.

#### External user contributions

The ModelSEED database, despite years of development, is still far from being perfect or complete. We welcome contributions from external users, including: (i) new proposed compounds, reactions, and pathways; (ii) curations to existing data including correcting reaction stoichiometry, aliases, or molecular structures; and/or (iii) integrating new tools to support database maintenance, quality control, and analysis. Users can propose changes by creating their own fork of the ModelSEED GitHub repository, implementing their changes within this fork, running ModelSEED test scripts to ensure that the proposed changes meet data quality and minimal information standards, and submitting changes to the ModelSEED team for review by issuing a pull request in GitHub against the dev branch. Once the team has ensured that proposed changes meet all standards, pull requests can be merged. The pull request mechanism on GitHub includes a built-in discussion forum to permit interactive discussion of proposed changes. We utilize Travis CI (56) along with scripts for testing data immediately, and reporting whether or not data in the pull request is valid.

#### Release procedure

On a quarterly basis, we will release a new version of the ModelSEED database via GitHub. Releases will always be deployed from the master branch in GitHub, and each release will be tagged with a version in GitHub. Additionally, on release, updated data will be deployed into the ModelSEED modeling environment as well as the chemistry database in the U.S. Department of Energy (DOE) Systems Biology Knowledgebase, KBase (57).

## RESULTS

### Growth in the compounds and reactions included in the ModelSEED database

Development of the ModelSEED database began ten years ago with the release of the first ModelSEED resource for microbial metabolic model reconstruction (58). The database was expanded in 2014 by integrating additional sources of plant biochemistry with the release of the PlantSEED (59). Here, for the first time, we are releasing the ModelSEED biochemistry database as a stand-alone resource. This new release includes expansions of the ModelSEED, updating data from our source databases, and adding additional sources. As expected, the ModelSEED database has expanded over time from 13,257 reactions in 2010 to 44,031 reactions today (Table 2).

**Table 2.** The statistics of the ModelSEED biochemistry database over time (Figure 1). *A complete reaction is one where every reactant has a fully defined metabolic structure in our database.

|  | 2010 (ModelSEED) | 2014 (PlantSEED) | Current 2020 |
| --- | --- | --- | --- |
| **Compounds** | **16,275** | **27,694** | **33,995** |
| Compounds with Structures | 13,821 (85%) | 19,605 (71%) | 28,503 (84%) |
| Compounds with generic groups | 1,261 (8%) | 1,402 (5%) | 4,368 (13%) |
| **Reactions** | **13,257** | **27,558** | **36,645** |
| Complete* Reactions | 11,338 (86%) | 7,898 (29%) | 28,947 (79%) |
| Balanced Reactions | 10,263 (77%) | 17,264 (63%) | 25,894 (71%) |
| Reactions with generic groups | 1,988 (15%) | 2,939 (11%) | 9,554 (26%) |

The updated ModelSEED database now contains reactions from KEGG, MetaCyc, BiGG, MetaNetX, and Rhea, but it is important to note that only the KEGG and MetaCyc databases have been integrated in their entirety (Table 3).

**Table 3.** A description of the biochemistry data that we have integrated from various sources. Only KEGG and MetaCyc were completely integrated so the numbers for the other databases may not reflect their published content.

|  | ModelSEED | KEGG | MetaCyc | BiGG | MetaNetX | Rhea |
| --- | --- | --- | --- | --- | --- | --- |
| Compounds | 33,978 | 17,764 | 19,140 | 2,710 | 30,858 | - |
| Structures | 28,503 | 16,576 | 18,130 | - | - | - |
| Reactions | 36,645 | 10,861 | 22,097 | 4,337 | 23,757 | 8,870 |

Molecular structure is very important in the ModelSEED because we use structure as our primary tool to map together identical compounds from our source databases and because we apply structures with thermodynamic property estimation tools to predict Gibbs free energy change for compounds and reactions. Currently, 84% of compounds in the ModelSEED have specified structure, which translates to 81% of reactions defined as *Complete*, meaning the structure is defined for every reactant involved in the reaction (Table 2).

One significant area of improvement for this new release of the ModelSEED was the identification and correction of redundant copies of various compounds and reactions that were previously added to the database in error due to a failure to match identical compounds. We previously failed to match identical compounds based on three problems: (i) no associated molecular structure, making it impossible to automatically match these compounds to other compounds in our database based on structure; (ii) errors or inconsistencies in compound structures that prevented a match from being made; or (iii) missing stereochemistry information in compound structures. We identified and corrected some of these issues in this latest release by reviewing and correcting many problems. For example, we reviewed 47 cases where sets of two or more compounds in our database appeared to have completely identical structures, involving 3,320 reactions. Ultimately 32 compounds were consolidated, which led to the correction of 1,301 reactions in our database that involved one or more of these compounds as reactants. This subsequently led to the consolidation of 7,577 reactions. We were also able to identify and correct previously automated consolidations of compounds and reactions that turned out to be erroneous. These cases primarily consisted of stereochemically generic compounds being consolidated with stereochemically specific versions. In the majority of cases, we only had to ensure the correct structure was used. We corrected the remainder of cases by disambiguating 39 compounds, leading to the correction of 20 reactions. We note that this is an ongoing curation effort and our data is by no means completely “fixed” or perfect but a work in progress. This is why we made the decision to officially distribute the ModelSEED database via GitHub (see a detailed discussion later). GitHub enables users to easily clone our database in a form that can also be easily updated as the database improves overtime (as well as showing users detailed information about every aspect of the data that have changed).

### Thermodynamics

For this release of the ModelSEED database, eQuilibrator (54) was applied to update Δ_*f*_*G*′^*o*^ for the compounds in the database. Δ_*f*_*G*′^*o*^ could be computed by eQuilibrator for 17,602 (62%) of the compounds with assigned structures in the database. The remaining 10,885 compounds had structures representing abstract molecules, macromolecules, or in some cases compounds containing functional groups with no associated energy contribution in eQuilibrator. 9,962 (57%) of the successful Δ_*f*_*G*′^*o*^ predictions from eQuilibrator had very low uncertainty (<5%), but 6,939 (39%) had a high uncertainty of over 100%. Given the large disparity in uncertainty values between eQuilibrator and our previously applied group contribution method (53) for computing Δ_*f*_*G*′^*o*^, we retain the Δ_*f*_*G*′^*o*^ values from both methods in our repository, with the source and uncertainty of each value labeled.

We similarly applied eQuilibrator to predict new Δ_*r*_*G*′^*o*^ for the reactions in the ModelSEED database, where we observed similar results with 13,429 (84%) reactions having an uncertainty below 5% and 2,503 (16%) reactions having an uncertainty over 100%. As with the Δ_*f*_*G*′^*o*^ predictions, we retained both our original Δ_*r*_*G*′^*o*^ and the eQuilibrator Δ_*r*_*G*′^*o*^ values in our repository, with the source and uncertainty of each value labeled. Unlike with the Δ_*f*_*G*′^*o*^ values, it may be possible to mix and match Δ_*r*_*G*′^*o*^ values from these competing methods. The results from our recomputation of Δ_*f*_*G*′^*o*^ and Δ_*r*_*G*′^*o*^ using eQuilibrator and our original group contribution method are highlighted in Table 4.

**Table 4.** A description of how the thermodynamics data were integrated. The total number of reactions includes any reactions for which any reagent has associated thermodynamics data, and the number of complete reactions excludes reactions for which any reagent does not have associated thermodynamics data. GC = Group Contribution; eQ = eQuilibrator. *Accepted means the data were utilized if it passed several basic tests, but precedent was placed on data from eQuilibrator (see main text).

|  | Compounds | Reactions |
| --- | --- | --- |
| All | 33,978 | 36,645 |
| Structures (GC) | 28,487 (100%) | 36,645 (100%) |
| Complete (GC) | - | 19,486 (53%) |
| Accepted* (GC) | 9,267 (33%) | 6,307 (17%) |
| Structures (eQ) | 17,602 (62%) | 36,645 (100%) |
| Complete (eQ) | - | 17,763 (48%) |
| Accepted* (eQ) | 10,663 (37%) | 13,694 (37%) |
| Final | 19,930 (70%) | 20,001 (55%) |

When comparing the newly calculated Δ_*r*_*G*′^*o*^ computed by eQuilibrator to that computed using the group contribution method, we find that most (65%) of the reactions had a difference of less than 5 kcal/mol (Figure 2). As the difference increases, the number of reactions dropped significantly, only 9% of the reactions had a difference above 15 kcal/mol. The uncertainty of the Δ_*r*_*G*′^*o*^ computed by eQuilibrator for reactions with a difference greater than 15 kcal/mol was also high, at least 80% of these reactions had more than 100% uncertainty, which implies a low confidence in the eQuilibrator energies for these reactions with high deviations in their Δ_*r*_*G*′^*o*^ values. When directly comparing the uncertainty between the two approaches, 43% of the compared reactions exhibited more than 100% uncertainty for the group contribution method and only 10% of the compared reactions exhibited more than 100% uncertainty for eQuilibrator (Figure 2). For the purpose of this analysis We only compared non-transport and mass-balanced reactions

**Figure 2.**
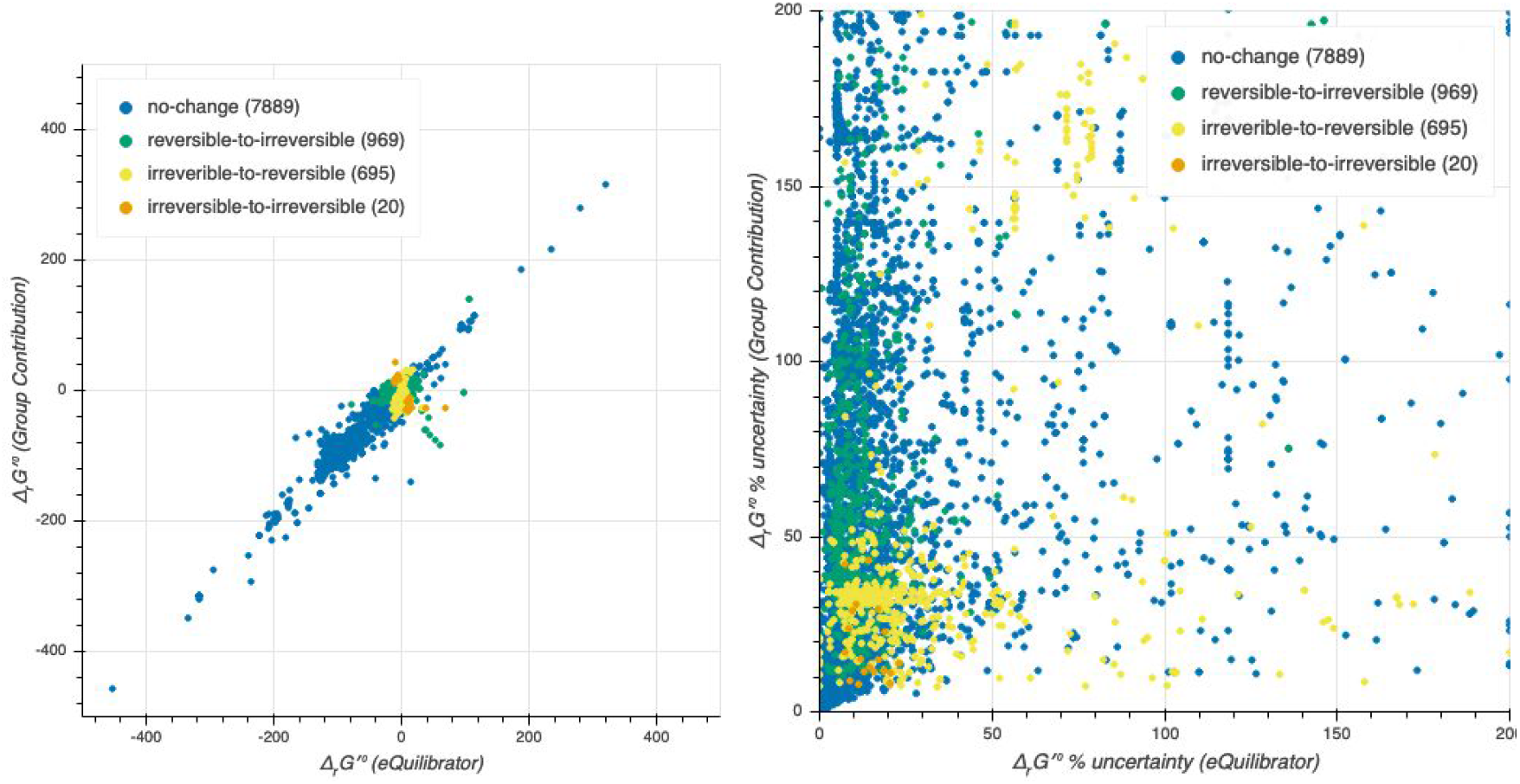
Thermodynamics in ModelSEED and eQuilibrator. We integrated the results calculated by eQuilibrator, a more recently developed approach, to estimate the gibbs energy of formation for more than 17K compounds and 36K reactions in the database. In comparing the results for reactions to those of the group contribution approach used in this and prior releases of ModelSEED, we find that there’s very little change (left panel), but in a small percentage of cases, the error reported for values computed by eQuilibrator is greater than 100% (right panel). The integration of data computed by eQuilibrator led to the adjustment of thermodynamic reversibility for roughly ~7% of the reactions in our database (see “Thermodynamics” section).

**Figure 3.**
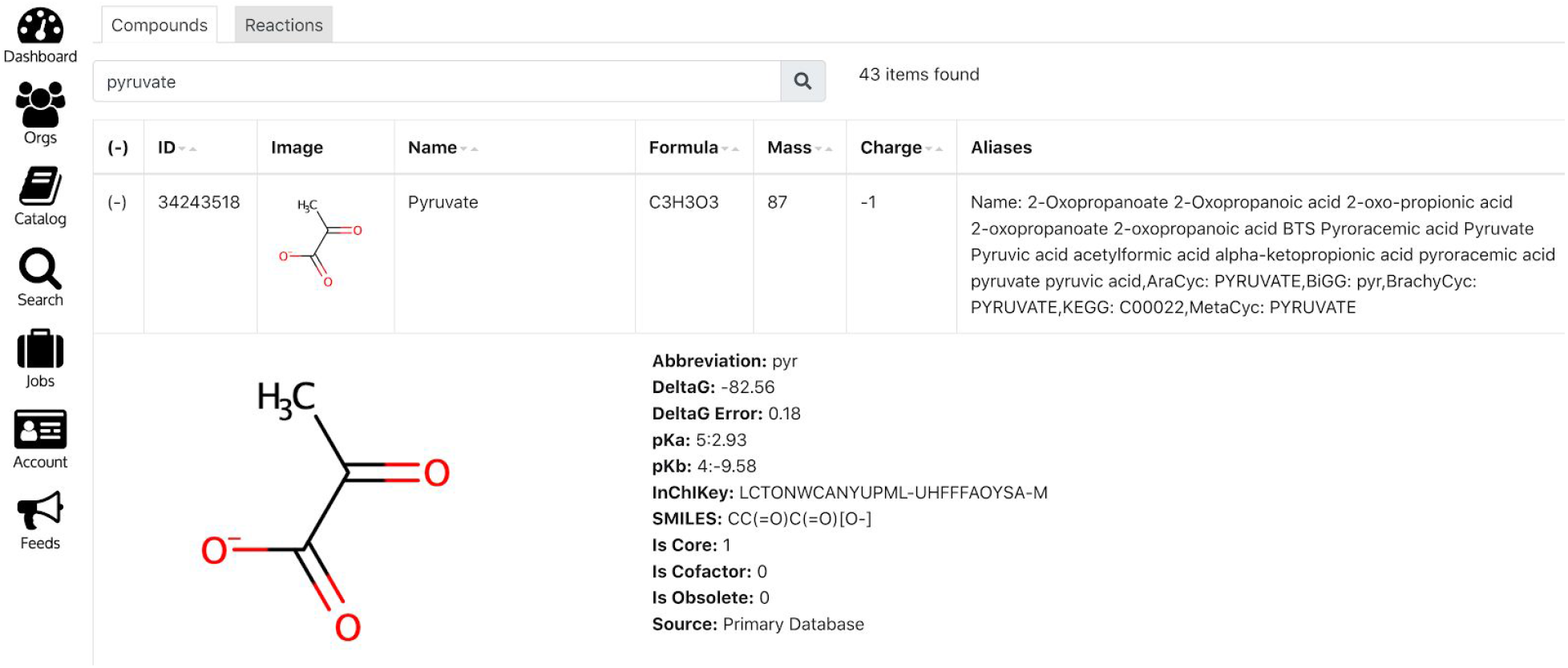
Biochemistry in DOE Systems Biology Knowledgebase (KBase). The ModelSEED biochemistry database is widely used for a range of metabolic modeling Apps in KBase Narratives (https://kbase.us) and there is also an interface for searching the entire biochemistry. The screenshot shows an example of a search result where “pyruvate” is used for the search term, and the first returned hit is expanded (by clicking in the first column).

We apply these newly calculated reaction energies, using the eQuilibrator values when the uncertainty was low and using the group contribution method values otherwise, to determine the reversibility of all reactions in the ModelSEED (see methods). Based on this analysis, we find that we were able to determine the reversibility of 4,118 reactions that had not been determined before. For the reactions for which reversibility had been determined before, we compared the reversibility if we were to use the group contribution method alone versus if we were to apply eQuilibrator values (Figure 2). We find that 82% of the compared reactions exhibited no change, while the remaining 18% did. Having adopted the approach integrating eQuilibrator values, we cross-checked these reactions with the conditional reactions in our templates to ensure that they did not disrupt the automated reconstructions in ModelSEED and PlantSEED, and we integrated the updated reversibility constraints into the gap-filling reactions available in the ModelSEED and PlantSEED templates.

### Improvements to database connectivity

One key purpose driving the development of the ModelSEED biochemistry database was to serve as the underlying chemistry source for the reconstruction of metabolic models in the ModelSEED (58), PlantSEED(58, 60), and KBase (57). As such, we needed to evaluate how our overall database performs in flux balance analysis. Metabolic modeling and flux balance analysis both place very specific demands on a biochemistry database. First and foremost, FBA can only be applied to reaction networks comprised of exclusively mass balanced reactions, so imbalanced reactions must be filtered out of any network that is to be leveraged for FBA. This does not mean that imbalanced reactions are excluded from the ModelSEED database, as it was also important to have as complete a database as possible to support annotation comparison, which is discussed later. However, all imbalanced reactions were identified and filtered from the database prior to the use of data in any FBA-based approach. Currently, of the 36K reactions in the ModelSEED database, 32,254 (73.3%) are charge and mass balanced, 2,548 (5.8%) are mass balanced but not charge balanced, and 9,229 (20.9%) are not mass balanced. This represents a significant improvement over previous releases which included only 22,421 mass balanced reactions.

Secondly, FBA also depends on having constraints on the reversibility of reactions to ensure that reactions have the flexibility needed to replicate true biological behavior without having too much flexibility leading to thermodynamically unrealistic flux profiles. For this, the improved Δ_*r*_*G*′^*o*^ estimates in this release of the ModelSEED lead to an adjustment of the reversibility rules associated with 2,783 reactions as described in the previous section. These changes impact the connectivity and overall feasibility of the ModelSEED reaction network when applied with flux balance analysis, as described.

We tested the impact of all of these changes on the connectivity and flux profile of our overall reaction network when used with flux balance analysis (Table 5). First, we applied flux balance analysis to determine how many reactions in our database were functional, meaning they were capable of carrying a nonzero mass-balanced flux from one set of transported metabolite inputs to another set of transported metabolite outputs (61). We actually saw a very slight decline in the number of functional reactions versus our previous release with the PlantSEED. This is primarily a result of our efforts to combine and eliminate redundant reactions in our database (described earlier). To test the extent to which this new release has improved capabilities in simulating phenotypes, we applied our database to simulate biomass production, using an example bacterial biomass objective function, in 390 Biolog growth conditions (62). In this study, our database was capable of successfully producing biomass for 353 (91%) of the Biolog conditions, which was an improvement over previous releases.

**Table 5.** Results from running flux balance analysis on ModelSEED database. A reaction is considered functional if it were determined to be capable of carrying nonzero mass-balanced flux (see main text). *Values in parentheses are without thermodynamic constraints.

| Release | Total reactions | Mass balanced | Reversible | Functional reactions* | Functional growth conditions |
| --- | --- | --- | --- | --- | --- |
| Current (2020) | 36,645 | 25,894 | 18,669 | 21,841<br>(23,029) | 353 (91%) |
| PlantSEED (2014) | 27,558 | 17,264 | 8,906 | 21,916<br>(23,460) | 337 (86%) |
| ModelSEED (2010) | 13,257 | 10,263 | 6,195 | 8,504<br>(9,072) | 330 (85%) |

### Leveraging ModelSEED biochemistry to map annotation ontologies and support comparison

The field of biology has long struggled to arrive at a standardized controlled way of describing the function of genes and their products. A variety of controlled vocabularies do exist (e.g. UniRef (63), Enzyme Classification numbers (64), Gene Ontology (65), KEGG orthology (48), SEED (66)), but many annotation platforms do not use these controlled vocabularies. Additionally, in order to compare the annotations of one platform with those of another, it is critically important to map together equivalent ontology terms. This enables one to differentiate cases where a difference in the function assigned to a gene by two platforms represents an actual disagreement in the function of the gene rather than a difference in nomenclature. Unfortunately, in the absence of any other abstraction, this mapping of functional descriptions becomes a largely manual exercise in syntactic interpretation (e.g., is the function described by these words equivalent to the function described by these other words). Fortunately, in the case of metabolic functions, we have another abstraction: biochemistry. Biochemistry is distinctive in that, if one knows the stoichiometry of a reaction and the molecular structure of the metabolites involved in the reaction, one does not need to rely on alias mapping or syntactic interpretation to determine if one reaction is equivalent to another. Instead, it is possible to computationally encode the compound structure and associated reactions into unique strings that make it possible to automatically compare reactions from different databases to one another. This makes biochemistry a valuable tool that may be leveraged to automatically map between functional ontologies where those ontologies have been associated with some kind of biochemistry database (which is increasingly becoming the case for most annotation ontologies).

Extensive effort has already been applied to exploit biochemistry structure codes (e.g. InChI and SMARTS) data to automatically generate translation tables among the reaction and compound identifiers in various biochemistry databases (e.g. MetRxn (67), MetaNetX (68)). Indeed, this semi-automated mapping is a major component of our own ongoing curation of the ModelSEED database, as previously discussed. We already highlighted one mechanism by which this automated mapping procedure may fail (e.g. if some compounds have missing or erroneous structures associated with them). The other significant reason why this mapping process may fail is that, in many cases, different biochemistry databases will represent the same biochemical pathway using different reactions (e.g. Figure 4b is an example of this in lipid metabolism). This issue can lead to significant apparent disagreement between the chemistry assigned by different resources to the same genes. For example, a comparison of the ModelSEED model of *E. coli* with the earliest manually curated model of *E. coli*, the iJR904 (69), revealed extensive apparent differences between these models. Of the 625 distinct compounds included in the iJR904 model, only 547 overlapped perfectly with the ModelSEED. However, many of these apparent miss-matching compounds were due to differences in representation of the same biochemistry between BiGG and ModelSEED.

**Figure 4.**
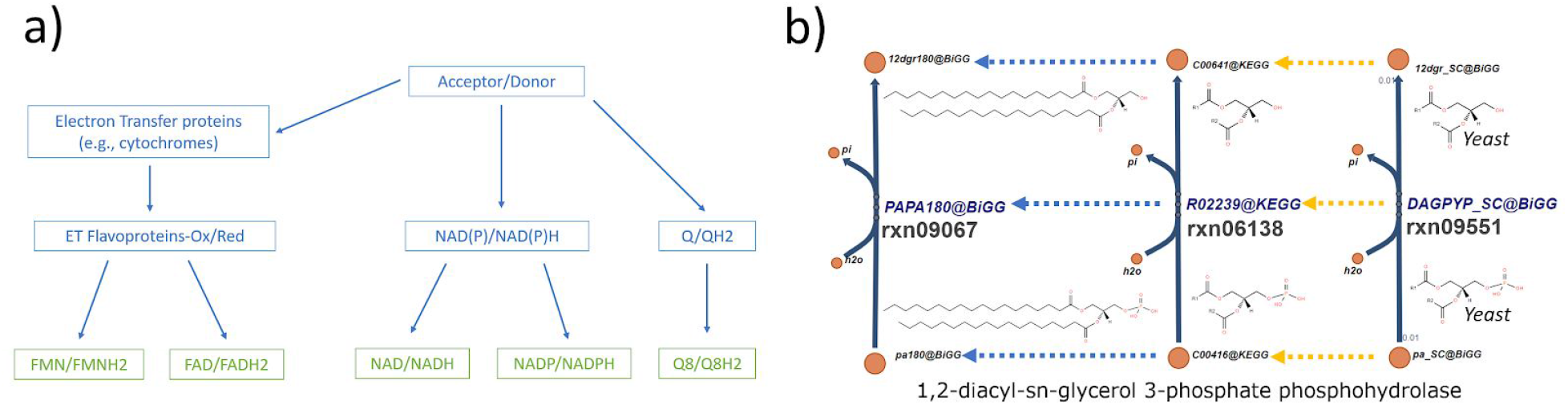
Directed acyclic graph representation of compound classes. Hierarchy is defined by their functional relationship in metabolism. a) Example of DAG representation of a few electron transfer compounds. b) Parallel representation of compound and reaction hierarchy; both rxn06138 and rxn09551 are abstract representations of rxn09067. However, rxn09551 is a context adapted version for the yeast, thus it has a different stoichiometry weight.

We have now developed a mechanism within the ModelSEED database for identifying and accounting for these differences in representation when comparing models and genome annotations. This approach begins with a general policy applied by the ModelSEED when we integrate chemistry from other resources into our own database. Unlike other biochemistry databases (e.g. KEGG, MetaCyc), the chemistry in the ModelSEED is not necessarily non-redundant. The same biochemistry may occur multiple times in the ModelSEED with differing representations. This is an explicit design decision in the ModelSEED made to facilitate the creation and maintenance of an ontological map between these various representations. The creation of this map begins with a semi-automated process of mapping together compounds in the database that are structurally different but chemically equivalent. Some associations can be made automatically (e.g. mapping **α**-D-Glucose and **β**-D-Glucose to D-Glucose and mapping D-Glucose and L-Glucose to generic Glucose). Note, other frameworks like ChEBI support this type of automated mapping as well (Hastings et al. 2009). Other associations must be made manually (e.g., mapping three different chain-length representations of fatty acid together as done in Figure 4b). Once these compound associations have been created, we then have an automated mechanism for the creation of a directed acyclic graph (DAG) connecting equivalent reactions to one another based on the associations among reactants. Once constructed, this DAG can be used to automatically support the translation of chemistry from one mapping to another, which in turn enables the automated comparison of metabolic annotations between resources. Applying our current DAG to our example comparison of the ModelSEED and iJR904 model of *E. coli*, this translation process reduced the number of mismatching compounds from 78 to 31. The impact of the reactions (Figure 5) is also significant, the number of uniques detected in iJR904 is reduced from 258 to 159, the usage of different isomers had the highest impact, followed by abstract representation of phospholipids and lumped fatty acid metabolism.

**Figure 5.**
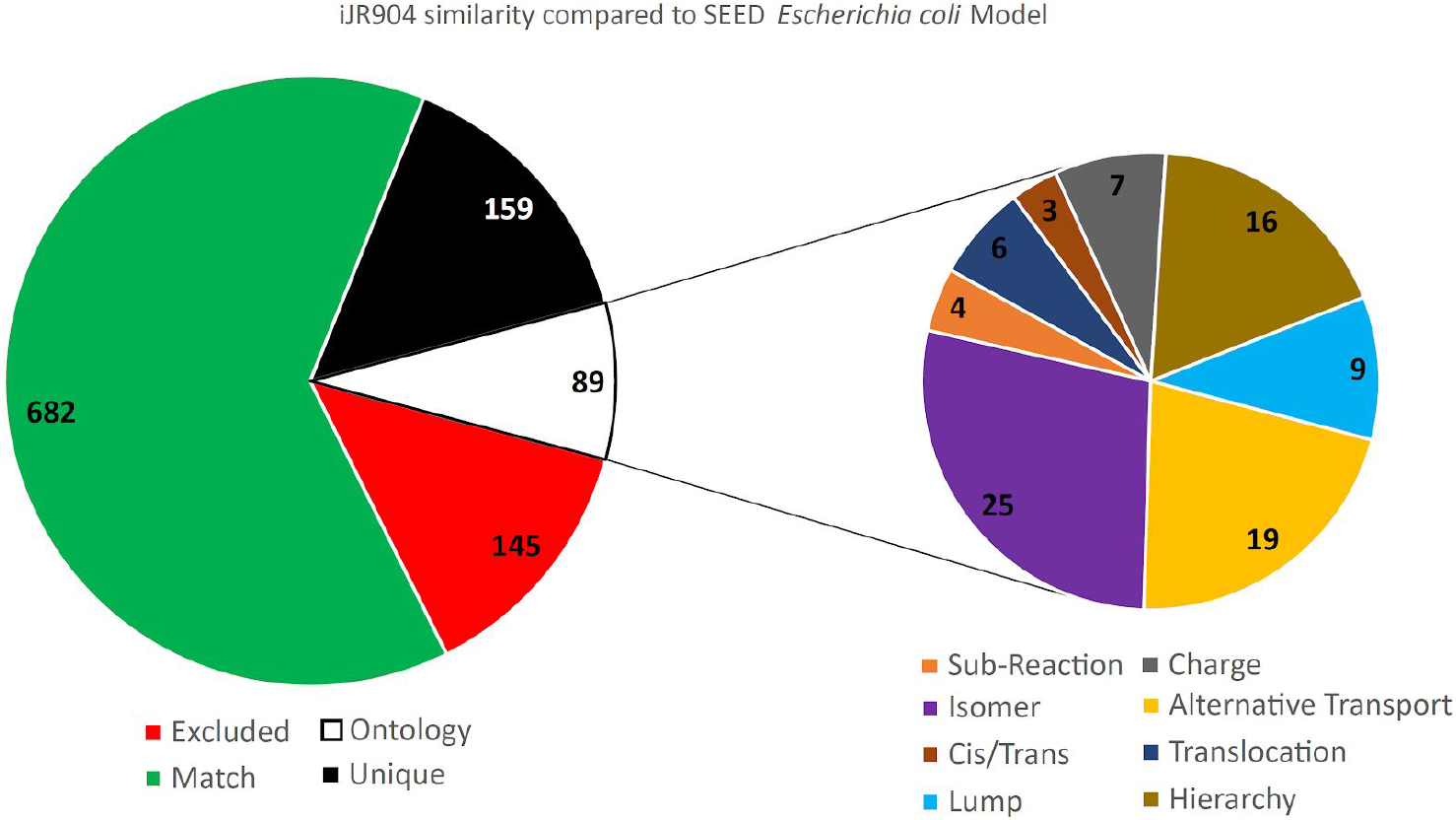
Reaction comparison between iJR904 and ModelSEED biochemistry of the *Escherichia coli* genome-scale model. Excluded - Exchange reactions, Biomass, ATPM; Match - Traditional matching approach (identity matching) with protonation comparison. Unique - Reactions that are not present in ModelSEED Model. Sub chart: Reactions otherwise marked as unique but with alternative representation in ModelSEED. Isomer and Cis/Trans - Reaction present but utilizing different isomer or cis/trans metabolite; Sub-Reaction/Lump - Reaction present but is a merge or split version of a ModelSEED reaction; Charge - Reaction matches same compound but with different charge; Hierarchy - Reaction matches ModelSEED reaction but utilizing an abstract representation of the compound; Translocation - Reaction matches exact stoichiometry but utilizing different compartment configuration; Alternative Transport - Transport of the compound present but using different mechanism or co-substrate. The submitted manuscript has been created by UChicago Argonne, LLC as Operator of Argonne National Laboratory (“Argonne”) under Contract No. DE-AC02-06CH11357 with the U.S. Department of Energy. The U.S. Government retains for itself, and others acting on its behalf, a paid-up, nonexclusive, irrevocable worldwide license in said article to reproduce, prepare derivative works, distribute copies to the public, and perform publicly and display publicly, by or on behalf of the Government. The Department of Energy will provide public access to these results of federally sponsored research in accordance with the DOE Public Access Plan.

The current ontological mappings established in the ModelSEED represent three different types of relationships: (1) equivalent compound sets; (2) lumped reaction sets; and (3) context-specific reaction sets. We stored each of these sets in separate DAGs that connect together ModelSEED compound and reaction entities. The equivalent compound and reaction set relationships expose compounds and reactions that are abstract/generic representations of other compounds and reactions. For example, in many electron transfer reactions (Figure 4a) when the cofactor is unknown, both KEGG and MetaCyc use a pair of abstract compounds to act as placeholders (Acceptor/Donor). These reactions must be adapted when used for modelling purposes. KEGG and MetaCyc currently contain 235 and 644 reactions respectively that are represented with generic acceptors. Using the ontological relationships implemented in the ModelSEED, we identified 104 clusters of reactions with specific cofactors that represent instantiations of generic acceptor/donor reactions. Now if two models from two different sources use different members of these clusters to represent the same overall reaction, we can automatically determine that these models at least agree that the generic reaction is happening. Other common problematic abstractions include: sugar isomers, and abstract representations of repetitive pathways (e.g., phospholipids, quinones, etc). The lumped reaction set relationship connects lumped versions of reactions to the series of sub-step reactions that have been lumped. We use these relationships to automatically convert lumped reactions into their component unlumped reactions or vice versa. The organism/context specific relationship connects entities that were modified to fit a certain context (e.g., adaptation for modeling purposes) to their standard representation (Figure 4b). In general, these reactions are unfit for modelling purposes, but they might contribute knowledge for the database (e.g., related genes).

We note that the ModelSEED is far from the first biochemistry database to apply ontologies to metabolites and biochemical reactions. CheBi, MetaCyc, and KEGG all do this to varying extents (70). However, other database ontologies have focused on classifying and categorizing these entities, whereas our focus is on mapping data from disparate sources to better facilitate comparison, reconciliation, and integration of annotation information.

### Using GitHub as a tool for community contributions and maintenance

One major goal for this release of the ModelSEED biochemistry database is to become a community-driven resource, meaning that contributions, updates, and corrections could be rapidly integrated from the research community. We also want the database to evolve over time in as transparent a manner as possible. To accomplish these goals, we have released the ModelSEED database to the public, using the Creative Commons Attribution License, in the Git repository: https://github.com/ModelSEED/ModelSEEDDatabase. Note, data directly derived from KEGG and MetaCyc is still subject to licenses from these resources. Using Git to store the changes made to the data and to the underlying scripts inherently maintains provenance of the data and scripts.

All the main datasets in the repository are well-formatted, and accompanied by instructions and a library of scripts for loading and handling the data across several folders. The main compound and reaction databases were formatted as tables that can be exported to Excel and as structured JSON objects that can be directly imported into any scripting language and web application. For example, a researcher can load the data directly into a local Solr instance (as described below). These files are accompanied by the data and scripts we use to maintain metabolic structure, thermodynamics, and external identifiers and synonyms.

Crucially, we expect the ModelSEED database to grow and to be improved over time, and invite researchers to collaborate with us. The use of a Git repository in GitHub provides the means by which we can interact with researchers and include changes from external teams with the accompanying provenance. Researchers will be able to submit edits, additions, and changes to the current data via use of Git and GitHub Pull Requests. We particularly welcome any new metabolic pathways relevant to the microbial, fungal, and plant kingdoms, as well as any new metabolic structures or thermodynamic data that would improve the process of reconstructing metabolism. We will review these submissions, and interact with the wider community to merge the new data and maintain the repository at a high standard. Policies for community contributions to the ModelSEED Github are described in the methods. Finally, we will release new changes and data on a quarterly basis.

## DISCUSSION

Currently in the field of bioinformatics, there are many powerful techniques for predicting gene function, including numerous homology methods like BLAST, Hidden Markov Models, and k-mer indexing. There are also numerous non-homology methods exploiting chromosomal context, coexpression, gene fitness data, co-occurrence, and protein structure. No single approach is a panacea. Rather, it has been demonstrated numerous times that optimal results in bioinformatics are obtained by combining many different approaches and data sources together to obtain a consensus result (71). One of the biggest impediments to building such a consensus approach for biology today is the lack of a single standard ontology for describing gene functions. Another impediment is the need for a mechanism to be able to test predicting gene functions for consistency with available phenotypic evidence (e.g. growth conditions and gene fitness data). A final impediment is the need for a streamlined mechanism for the research community to rapidly integrate new annotations and pathways into these chemistry databases, as well as track full provenance on changes in those databases over time.

The ModelSEED biochemistry database was designed to address these challenges. By integrating together diverse chemistry databases, and building and maintaining mappings to those databases based on structure and ontology, we provide a resource that can automatically translate many different annotation ontologies into a single chemical representation. This in turn facilitates the rapid comparison and reconciliation of annotations. By also making that single chemical representation “modelable” and integrating deeply with model reconstruction platforms like the ModelSEED, PlantSEED, and KBase, we offer a means of converting that single chemical representation into mechanistic models. Mechanistic models can then apply functional annotations to predict conditional phenotypes like gene fitness or growth conditions so that competing annotations may be tested and reconciled to maximize consistency between phenotype predictions and experimental data. Finally, by deploying our database on GitHub, we provide an easy, trackable method for rapidly accepting contributions of new chemistry and data from the research community. GitHub also provides an excellent built in system for tracking database changes over time, as well as tracking who is responsible for each change.

Competing biochemistry resources do exist that meet one of these challenges. MetaCyc and BIGG are both top tier resources for supporting metabolic model reconstruction, but neither of these databases supports direct community contributions, and of this pair, only MetaCyc offers significant ontology support. Even in MetaCyc, the ontology support is directed more at classification rather than mapping between databases. Other resources exist that focus more specifically on supporting database mapping and ontology, including Rhea, which is integrated with gene ontology, and MetaNetX, which maintains mappings of identical compounds and reactions from numerous data sources. However, again neither of these resources supports direct community contribution or model reconstruction.

Mapping, reconciling, testing, and integrating knowledge of gene function in biological systems is one of the primary driving missions of KBase. The ModelSEED biochemistry database is an important part of the KBase platform. The tools presented here significantly advance that mission by providing a structured, extensible framework, with provenance, to support all of these activities for metabolism. As implemented in KBase, the ModelSEED biochemistry database exemplifies how a specialized, independently curated resource that provides valuable integration of multi-dimensional omics data can significantly enrich the available data content and structure in KBase, and thereby further empower a systems biological analytical approach for all KBase users.

## AVAILABILITY

We have released the ModelSEED biochemistry database to the public, using the Creative Commons Attribution License, in the Git repository: https://github.com/ModelSEED/ModelSEEDDatabase. The release of data will not only be in the repository but also deployed to several key resources: ModelSEED (https://modelseed.org) and its accompanying SOLR database (https://modelseed.org/solr) and KBase (https://kbase.us) by way of inclusion in all of our metabolic modeling Apps in KBase narratives, and also via the KBase search interface (https://narrative.kbase.us/#biochem-search).

In addition to our establishment of a GitHub repository for the ModelSEED data, we have also created a web interface in both the ModelSEED and KBase environments to search and browse this data. The ModelSEED interface to the biochemistry data is available at http://modelseed.org/biochem/reactions. This interface includes a compound and reaction table, fully searchable by column, including supporting search by aliases from other databases. The interface also includes compound and reaction landing pages showing a more detailed view of these entities. We also updated this interface to ensure that all biochemistry data are accessible without logging in. Finally, we added an interface to KBase for browsing this biochemistry data: https://narrative.kbase.us/#biochem-search. Like the ModelSEED interface, this tabular view enables users to search for reactions and compounds by a variety of terms, including aliases, and redirects compound and reaction views to the landing pages in the ModelSEED.

In addition to these web interfaces for manually browsing the ModelSEED biochemistry data, we have also created programmatic APIs. All data can be loaded into an Apache Solr (https://lucene.apache.org/solr/) database, which offers a publically accessible REST API for accessing the data. In the Solr folder of the ModelSEED GitHub, we have several examples of how a researcher can fetch the data directly via https, or via a python script. If researchers wish to set-up their own Solr endpoint to serve their own biochemistry data, formatted in the same manner as our public one, a set of instructions for how to do that is in the same folder.

## SUPPLEMENTARY DATA

Supplementary Data are available at NAR online.

