## Supplemental Table 1 for "The ModelSEED Database for the integration of metabolic annotations and the reconstruction, comparison, and analysis of metabolic models for plants, fungi, and microbes"

**Supporting Table S1:** Sources of biochemistry integrated into PlantSEED

| Database/Model | Species | Owner/Reference | Date / Version | Number of Compounds | Number of Reactions |
|---|---|---|---|---|---|
| KEGG | N/A | KEGG (1, 2) | 12/03/18 | 17,793 | 14,039 |
| MetaCyc | N/A | PathwayTools (3) | 22.5 | 19,172 | 27,414 |
| EcoCyc | *Escherichia coli* | PathwayTools (3) | 16.1 | 3,355 | 2,411 |
| PlantCyc | N/A | PMN (4, 5) | 7.0 | 3,919 | 4,436 |
| AraCyc | *Arabidopsis thaliana* | PMN (5) | 10.0 | 4,229 | 4,008 |
| PoplarCyc | *Populus trichocarpa* | PMN (4) | 4.0 | 2,318 | 2,301 |
| SoyCyc | *Glycine max* | PMN | 2.0 | 2,796 | 2,657 |
| ChlamyCyc | *Chlamydomonas reinhardtii* | PMN | 03/03/10 | 1,718 | 1,875 |
| BrachyCyc | *Brachypodium distachyon* | Gramene (6, 7) | 2.0 | 2,348 | 3,104 |
| MaizeCyc | *Zea mays* | Gramene (8) | 2.0 | 2,219 | 2,358 |
| RiceCyc | *Oryza sativa* | Gramene (9) | 2.0.1 | 1,859 | 1,955 |
| SorghumCyc | *Sorghum bicolor* | Gramene (6, 7) | 1.0.1 | 1,773 | 1,873 |
| iAF1260 | *Escherichia coli* | Feist *et al.* (2007) (10) | | 1,041 | 2,064 |
| iAF692 | *Methanosarcina barkeri* | Feist *et al.* (2006) (11) | | 562 | 613 |
| iIN800 | *Saccharomyces cereivisae* | Nookaew *et al.* (2008) (12) | | 683 | 1,053 |
| iJR904 | *Escherichia coli* | Reed *et al.* (2003) (13) | | 629 | 921 |
| iMA945 | *Salmonella spp.* | AbuOun *et al.* (2009) (14) | | 1,032 | 1,960 |
| iMM904 | *Saccharomyces cerevisiae* | Mo *et al.* (2009) (15) | | 712 | 1,401 |
| iRR1083 | *Salmonella typhimurium* LT2 | Raghunathan *et al.* (2009) (16) | | 759 | 1,086 |
| iSB619 | *Staphylococcus aureus* N315 | Becker *et al.* (2005) (17) | | 614 | 639 |
| iSO783 | *Shewanella oneidensis* MR-1 | Pinchuk *et al.* (2010) (18) | | 634 | 774 |
| iAbaylyiv4 | *Acinetobacter baylyi* ADP1 | Durot *et al.* (2008) (19) | | 699 | 867 |
| * | *Bacillus subtilis* | Goelzer *et al.* (2008) (20) | | 475 | 504 |
| iGT196 | *Buchnera aphidicola* | Thomas *et al.* (2009) (21) | | 740 | 210 |
| iIT341 | *Helicobacter pylori* | Thiele *et al.* (2005) (22) | | 411 | 473 |
| iJN746 | *Pseudomonas putida* KT2440 | Nogales *et al.* (2008) (23) | | 706 | 915 |
| iMO1056 | *Pseudomonas aeruginosa* PAO1 | Oberhardt *et al.* (2008) (24) | | 750 | 864 |
| iND750 | *Saccharomyces cerevisiae* | Duarte *et al.* (2004) (25) | | 650 | 1,038 |

| | | | | | |
|---|---|---|---|---|---|
| iNJ661 | *Mycobacterium tuberculosis* H37Rv | Jamshidi & Palsson (2007) (26) | | 761 | 951 |
| iPS189 | *Mycoplasma genitalium* | Suthers *et al.* (2009) (27) | | 277 | 262 |
| iRS1563 | *Zea mays* | Saha *et al.* (2011) (28) | | 1,812 | 1,949 |
| iRS1597 | *Arabidopsis thaliana* | Saha *et al.* (2011) (28) | | 1,759 | 1,837 |
| iYO844 | *Bacillus subtilis* | Oh *et al.* (2007) (29) | | 776 | 1,016 |
| * | *Chlamydomonas reinhardtii* | Boyle & Morgan (2009) (30) | | 266 | 485 |
| * | *Chlamydomonas reinhardtii* | Manichaikul *et al.* (2009) (31) | | 124 | 238 |
| * | *Chlamydomonas reinhardtii* | Chang *et al.* (2011) (32) | | 1,164 | 2,084 |
| C4GEM | *Zea mays* | D'al Molin *et al.* (2010) (33) | | 1,207 | 1,227 |
| * | *Arabidopsis thaliana* | Mintz-Oron *et al.* (2012) (34) | | 1,181 | 3,382 |
| * | *Arabidopsis thaliana* | Poolman *et al.* (2009) (35) | | 1,224 | 1,354 |
| AraGEM | *Arabidopsis thaliana* | D'al Molin *et al.* (2010) (36) | | 1,546 | 1,590 |
| AlgaGEM | *Chlamydomonas reinhardtii* | D'al Molin *et al.* (2011) (37) | | 1,662 | 1,713 |

The numbers of compounds and reactions are listed for each source *after* integration, and may not reflect the numbers seen in the literature. In addition, the number of reactions includes compartmentalized reactions, which may be duplicates. *No unique identifier for these models was described in the literature.
